# PDP-Miner: an AI/ML tool to detect prophage tail proteins with depolymerase domains across thousands of bacterial genomes

**DOI:** 10.1101/2025.01.20.633936

**Authors:** Jeff Gauthier, Irena Kukavica-Ibrulj, Roger C. Levesque

## Abstract

**Motivation:** Antibiotic resistance is predicted to become the leading cause of human mortality by 2050. Despite this, no other major antibiotic class has been approved for medical use since 1987. Nevertheless, phage tail proteins offer a promising alternative, given their depolymerase activity toward outer membrane polysaccharides. Several pathogenic bacteria harbor prophages, thus making these prophages’ molecular target already known.

**Results:** We therefore developed a wrapper for an existing machine learning-based phage depolymerase prediction tool (Depolymerase-Predictor), called PDP-Miner, which annotates phage tail proteins *ab initio*, detects depolymerase activity within this candidate protein subset, and then performs post-hoc validation by annotating protein domains thereby allowing the user to investigate for protein domains indicative of depolymerase activity. This tool allowed identification of 10 high confidence phage depolymerase gene candidates across all 1,294 *Pseudomonas* genomes available on the International Pseudomonas Consortium Database and could likely help detecting other candidates across other genome databases as well.

**Availability and Implementation:** Source code is freely available for download at http:///www.github.com/jeffgauthier/pdpminer. Implemented in Bash, supported on native Linux or WSL and supports submitting subtasks to a SLURM workload queue. Requires Miniconda3 to install dependencies. This software is free and open source under the GNU General Public License v3.0.

**Supplementary information:** [add Supp. Mat. URL here when available]

## Introduction

Antibiotic resistance is predicted to become the leading cause of human mortality by 2050, surpassing cancer and cardiovascular disease (Antimicrobial Resistance Collaborators 2022). Despite this, no other major antibiotic class has been approved for medical use since 1987 (Hutchings, Truman and Wilkinson 2019), thereby contributing to the surge of AMR. There is an urgent need for novel, alternative antibiotics classes to mitigate AMR. Although penicillin, the first mass-produced antibiotic, was discovered by Fleming in 1929 (Fleming 1929), bacteriophages have been used as antibacterial treatments as early as in 1919 (Barron 2022). Phage therapy remains used to this day in Eastern European countries (Yang et al. 2023). However, the low reproducibility of manufacturing methods and their popularity in 20^th^-century Soviet republics may have contributed to their early dismissal in Western medicine in favor of broad-spectrum antimicrobial compounds (Loc-Carrillo and Abedon 2011).

Perhaps the most important caveat with phages is their narrow host range, often down to the strain level (Fong et al. 2021). Indeed, bacteriophages adhere to the cell wall via tail proteins (Taslem Mourosi et al. 2022) via highly specific binding to either: outer membrane receptors, lipopolysaccharides (LPS), exopolysaccharides (EPS), and capsule polysaccharides (CPS) or a combination of those. Several tail proteins also exhibit depolymerase activity toward specific types of these cell wall components to facilitate adsorption (Knecht, Veljkovic and Fieseler 2020). Therefore, using phage depolymerases (PDPs) alone (independently of the phage) to weaken the bacterial cell wall constitutes a potential treatment strategy (alone or in combination with antibiotics), assuming the PDP’s target is well known. This strategy has already delivered promising results in more than 20 animal infection model trials (Wang et al. 2024) (see Table 1). However, most PDPs currently described were purified from lytic phages (REF) which requires continuous culture to either maintain the phage or find its target.

**Table 1.** Candidate phage tail depolymerases found by PDP-Miner with DePP score above 75% and Pfam domains indicating depolymerase activity, across the IPCD genome collection (N = 1,294).

| IPCD isolate | Gene | contig | start | stop | frame | Pharokka annotation | Pfam domains | DePP score |
| --- | --- | --- | --- | --- | --- | --- | --- | --- |
| IPC_1377 | CDS_5089 | scaffold7.1 | 181786 | 183660 | + | tail protein | PF22432.1:PA2794-like_C | 92% |
| IPC_1419 | CDS_6000 | scaffold22.1 | 41837 | 43900 | + | tail protein | PF22432.1:PA2794-like_C | 88% |
| IPC_38 | CDS_4033 | scaffold7.1 | 99963 | 101723 | + | lytic tail protein | PF01464.25:SLT; PF01476.25:LysM | 77% |
| IPC_133 | CDS_2732 | scaffold5.1 | 323075 | 321303 | - | lytic tail protein | PF01464.25:SLT; PF01476.25:LysM | 77% |
| IPC_1078 | CDS_1576 | scaffold3.1 | 411575 | 409803 | - | lytic tail protein | PF01464.25:SLT; PF01476.25:LysM | 77% |
| IPC_135 | CDS_3183 | scaffold6.1 | 323084 | 321312 | - | lytic tail protein | PF01464.25:SLT; PF01476.25:LysM | 76% |
| IPC_37 | CDS_3048 | scaffold4.1 | 141331 | 143091 | + | lytic tail protein | PF01464.25:SLT; PF01476.25:LysM | 76% |
| IPC_1607 | CDS_2631 | scaffold5.1 | 324168 | 322396 | - | lytic tail protein | PF01464.25:SLT; PF01476.25:LysM | 76% |
| IPC_1751 | CDS_3739 | scaffold6.1 | 323078 | 321306 | - | lytic tail protein | PF01464.25:SLT; PF01476.25:LysM | 75% |
| IPC_1592 | CDS_2758 | scaffold6.1 | 323035 | 321263 | - | lytic tail protein | PF01464.25:SLT; PF01476.25:LysM | 75% |

Several pathogenic bacteria, *e*.*g. Pseudomonas aeruginosa*, are known to harbor prophages (up to 7 per genome) (Johnson, Banerjee and Putonti 2022), implying that if the strain’s outer cell wall components are well characterized, any PDP encoded by these prophages would have its target already known because the phage must have infected the strain at least once to insert its genetic material into its host’s genome. The PDP can therefore be isolated, purified and administered as a bactericidal treatment to other strains having similar extracellular polysaccharide structure, all without having to cultivate the source phage and host. Alternative strategies to produce a prophage tail protein would be complete gene synthesis, plasmid cloning, and expression at high levels. This in turn reduces the likelihood of genetic alterations to either the PDP or its source organism. Prophage DNA can be annotated from the host bacterial genomic DNA sequence.

In this study, we have investigated the presence of PDPs across 1,294 *P. aeruginosa* genomes from the International Pseudomonas Consortium Database (Freschi et al. 2015) (http://ipcd.ibis.ulaval.ca). Initially, the SVM-based tool Depolymerase-Predictor (DePP) was used alone because of this training set comprised of experimentally verified phage tail depolymerases (Magill and Skvortsov 2023), but it yielded several false-positive results when all protein-coding genes were considered as input. We therefore developed a wrapper for this software, called PDP-Miner, which annotates tail proteins *ab initio* then runs DePP exclusively on this subset. PDP-Miner also performs post-hoc validation by annotating protein domains thereby allowing the user to investigate for protein domains indicative of glycosyl hydrolase activity, among others.

This discovery pipeline could help mitigating AMR in *P. aeruginosa*, which is a major widespread human opportunistic pathogen (Weimann et al. 2024) infecting patients with cystic fibrosis (CF), chronic obstructive pulmonary disease (COPD), and large burn wounds, all while causing other soft tissue and systemic infections (Morin et al.). *P. aeruginosa*, a WHO top priority pathogen regarding AMR emergence (WHO 2024), accounts for half a million annual deaths (Ikuta et al. 2022).

## Materials and Methods

### Type strain dataset (4 genomes)

Genomic FASTA sequences from *P. aeruginosa* strains PAO1, PAK, LESB58 and *P. paraeruginosa* strain PA7 were retrieved from the Pseudomonas Genome Database (http://www.pseudomonas.com) to form a small dataset of phenotypically and genomically well-described strains in which PDP gene candidates are expected to be discovered. Indeed, PDP activity has been experimentally verified against these four type *Pseudomonas* strains in PAO1 (Olszak et al. 2017), PAK (Bhattacharjee et al. 2022), as well as the presence of phage-tail-like bacteriocins in PA7 (Saha et al.) and recently acquired prophage islands in Liverpool Epidemic Strain LESB58 (Winstanley et al. 2009).

### IPCD *Pseudomonas* genome dataset

All 1,294 *Pseudomonas* genomes available on the International Pseudomonas Consortium Database (IPCD, (Freschi et al. 2015) were selected and run through PDP-Miner. Beyond archiving source isolates, this database also routinely generated Illumina MiSeq 2×300 bp genome data assembled with the A5-miseq pipeline (Coil, Jospin and Darling 2015) for all isolates. Therefore, any potential PDP candidates found via PDP-Miner could be tested against the original strains and/or strains having a similar surface polysaccharide composition.

### PDP-Miner algorithm

An overview of the PDP-Miner algorithm, and how it was used in this study, are provided in **Figure 1**. Briefly, genome FASTA sequences are annotated with Pharokka v1.7.4 (Bouras et al. 2023) to list only phage-related genes. Then, phage tail protein genes are selected with seqtk v1.4 (https://github.com/lh3/seqtk), more precisely the “subseq” command (with list of loci labeled “tail protein” by Pharokka). Then, Depolymerase-Predictor v1.0.0 (DePP, (Magill and Skvortsov 2023), a SVM-based phage depolymerase classifier, assigns a probability score to only phage tail protein genes (i.e. their amino acid translation). DePP returns a 2-column table with gene IDs and a probability score. Pharokka and DePP annotations are then merged into a single table and only gene candidates with a DePP probability over 75% are kept. To further assess the validity of DePP predictions, PfamScan v1.6 (Mistry et al. 2021) is run on candidate gene products with over 75% DePP score. Domains found by PfamScan are inserted into the Pharokka/DePP table to allow the user to search for domains indicative of depolymerase activity, e.g. hydrolase (EC 3.1.-.-), lyase (EC 3.2.-.-), carbohydrate binding domains, phage tail domains, and more. This allows the user to judge and prioritize annotations by validating DePP’s prediction, which is known by us to generate false positives when non-phage genes are considered (Table S1).

**Figure 1.**
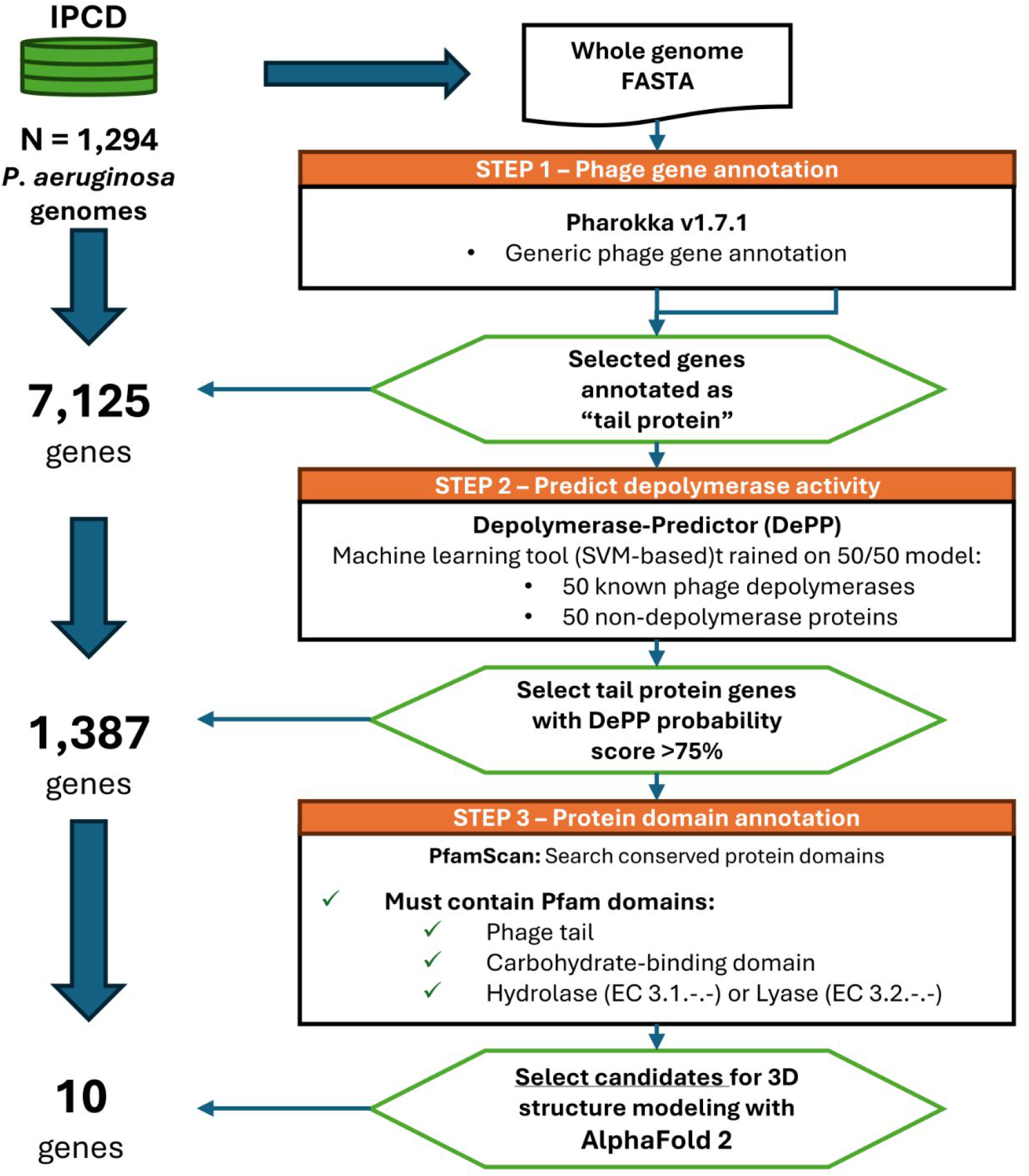
Flowchart describing the PDP-Miner pipeline. Briefly, prophage genes are annotated in silico with Pharokka (Bouras et al. 2023). Then, tail protein genes are scanned with Depolymerase-Predictor (DePP, Magill and Skortsov 2023) to obtain a probability score associated with each annotation. Finally, tail spike protein genes with >75% DePP probability are investigated for potential domains indicative of glycosyl-hydrolase/lyase activity with PfamScan (Mistry et al. 2020). Those candidate tail protein depolymerase genes are then selected for homology modeling with AlphaFold 2 (Jumper et al. 2021 Nature).

### Code and data availability

All IPCD genome sequences used in this study are publicly available through the NCBI BioProject primary accession code PRJNA325248. Raw sequencing data and strain metadata are available upon request at http://ipcd.ibis.ulaval.ca. The current implementation of PDP-Miner is publicly available on GitHub in the repository: http:///www.github.com/jeffgauthier/pdpminer,

## Results

### Initial type strain dataset (4 genomes)

The initial stage of the PDP-Miner workflow is to annotate phage tail proteins with Pharokka and evaluate their likelihood of being a depolymerase with Depolymerase-Predictor (DePP). This was initially tested with a restricted set of four *Pseudomonas spp*. type strain genomes (LESB58, PAK, PAO1 and *P. paraeruginosa* PA7). Among those, five putative phage tail proteins annotated by Pharokka were found with a DePP score above 75% (**Figure 2**). More precisely, two candidates were found for PA7 and one was found for each other strain. For these five candidate PDPs, the DePP score, *i*.*e*. the likelihood of them having depolymerase activity, ranged between 86% and 94%. In addition, we found by multiple sequence alignment that the nearest protein among DePP’s experimentally verified protein training set was the tail fiber protein of Klebsiella phage GH-K3 (**Suppl. Figure 1**), suggesting high structural similarity between the top 5 and GH-K3’s tail fiber protein.

**Figure 2.**
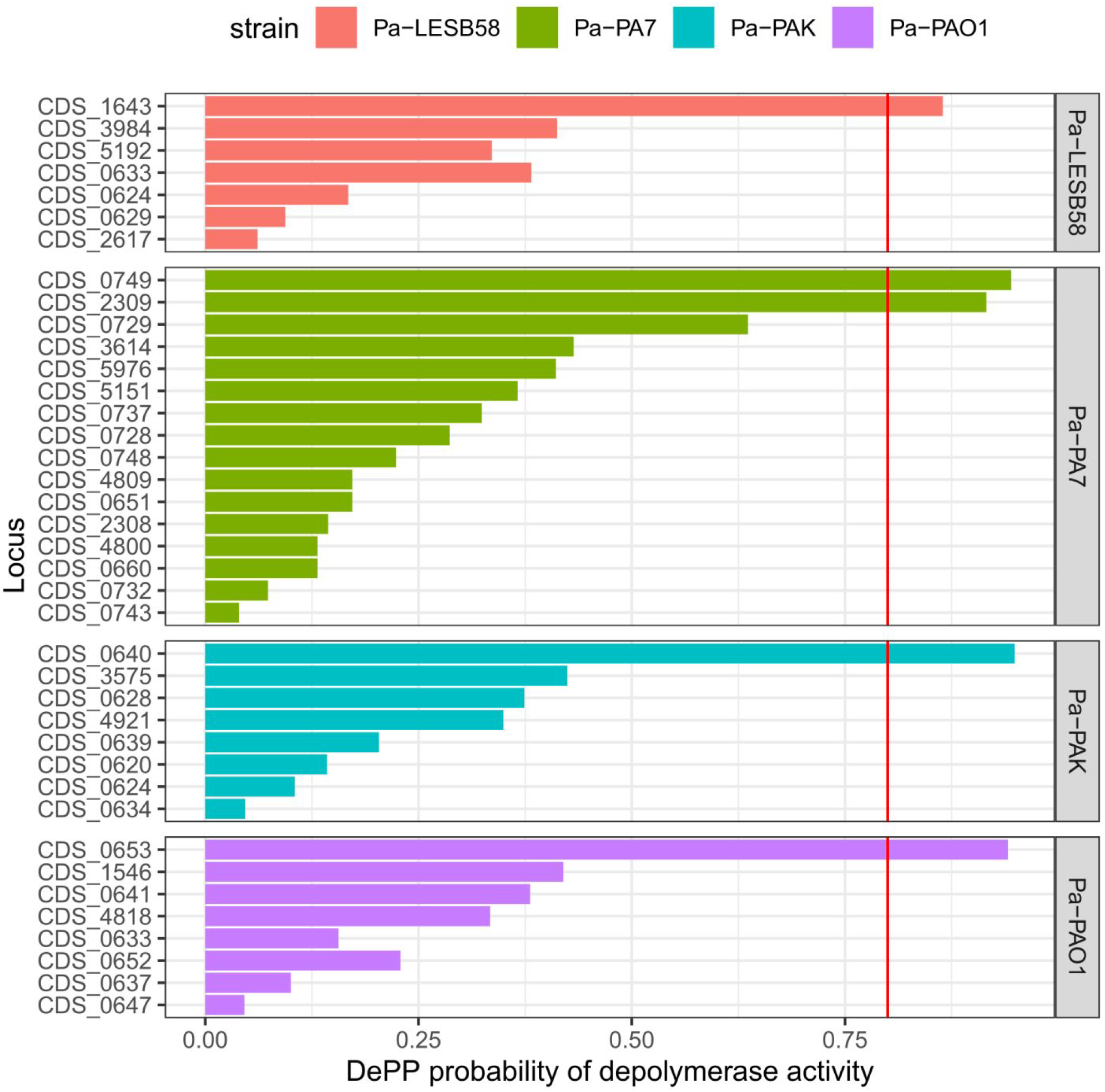
Phage tail proteins among four type Pseudomonas aeruginosa genomes PAO1, LESB58, PAK and PA7, sorted by DePP score. CDS: coding DNA sequence. DePP: Depolymerase-Predictor. The red vertical length indicates the user-specified threshold for selecting candidate phage depolymerases deserving further characterization at the domain level.

However, DePP by itself only reports a two-column table with sequence IDs and probability scores, which limits the interpretation of this dataset. To further help interpret the results, PDP-Miner uses PfamScan to annotate protein domains and report them alongside DePP score. Interestingly, phage-tail-3 and DUF1983 domains were found in GH-K3’s tail fiber protein and in each of the top 5 candidates, except the one in LESB58 which had only a phage-tail-3 domain (**Figure 3**). One protein, CDS #2309 from PA7, also had a “CBM_5_12” domain (Pfam domain 02839), which is a “carbohydrate binding domain found in many glycosyl hydrolase enzymes” (NCBI 2024).

**Figure 3.**
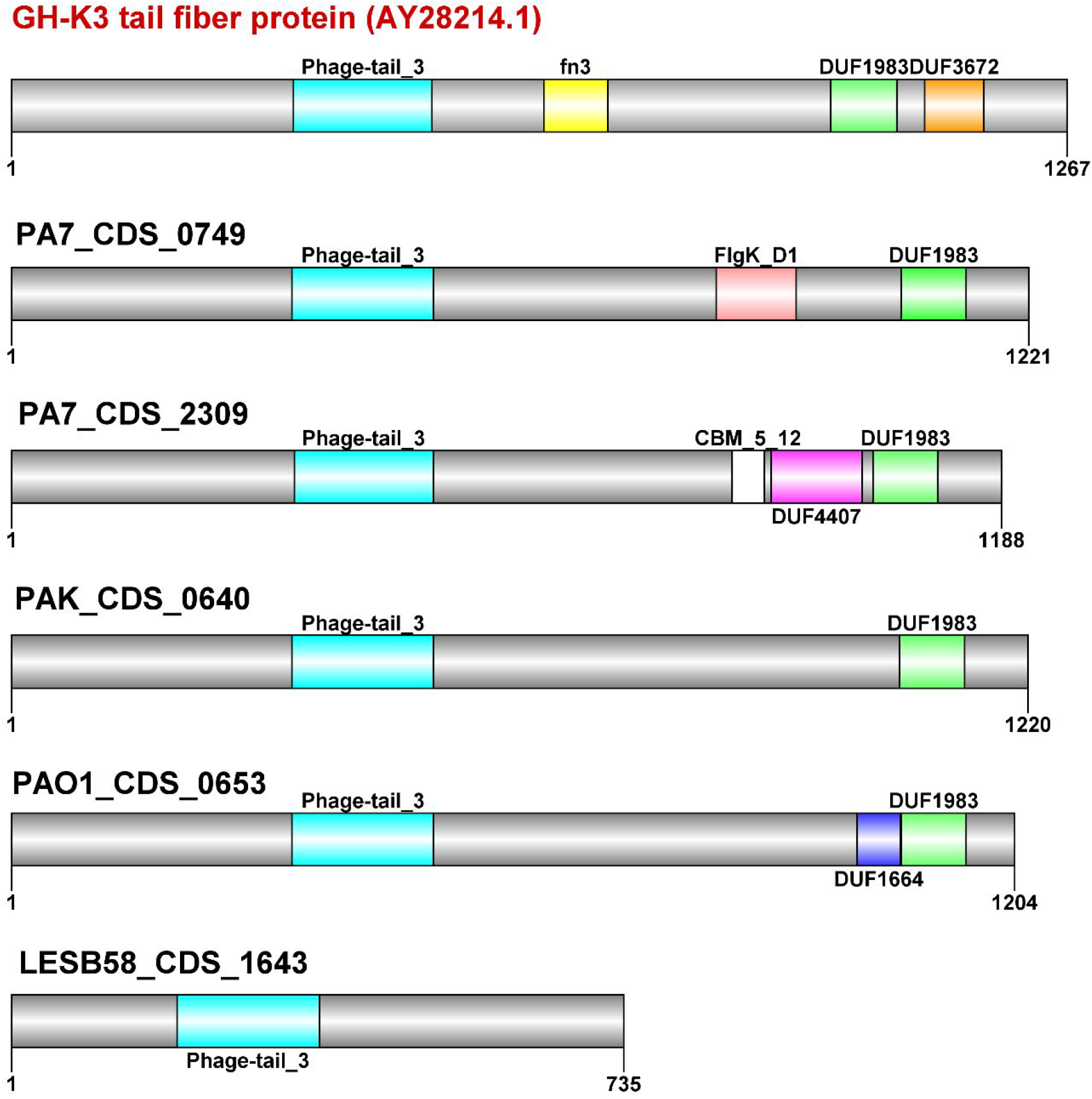
Protein domains of top scoring proteins among type Pseudomonas aeruginosa and paraeruginosa strains. All domain annotations follow Pfam nomenclature (fn3: fibronectin III, CBM: carbodydrate binding domain, flgK: flagellin K, DUF: domain of unknown function).

To further verify this structural similarity between GH-K3 and PA7 CDS #2309, and thereby its likelihood of being a phage tail depolymerase, we predicted their 3-dimensional structure with AlphaFold 2 via the NeuroSnap webserver (https://neurosnap.ai). The 3D structure of Depo32, a phage depolymerase, was used for guiding predictions as this structure was obtained through cryogenic electron microscopy (see PDB structure #7VYV). PA7 CDS #2309 and GH-K3’s tail fiber protein had highly similar overlapping structures (**Figure 4**).

**Figure 4.**
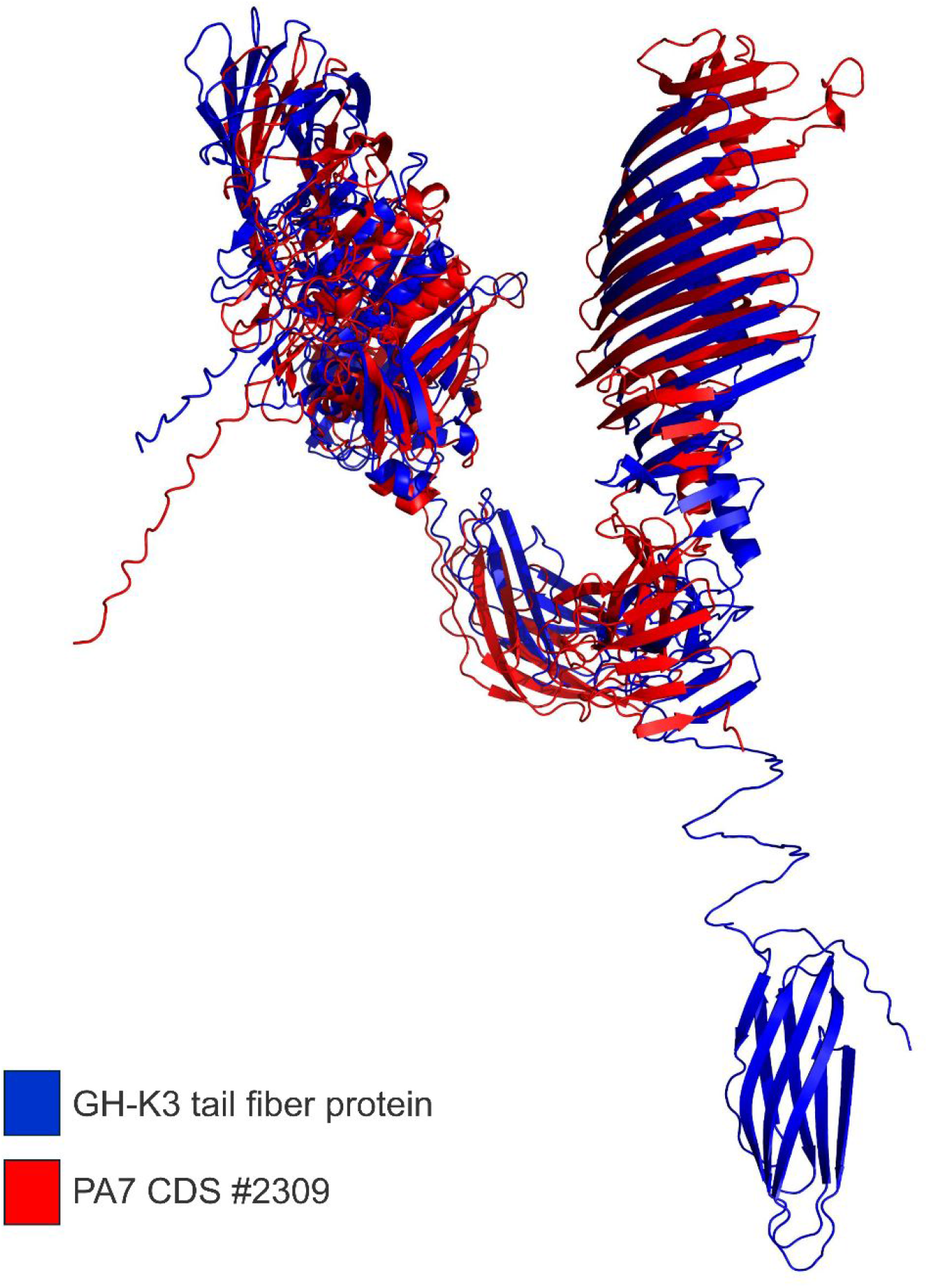
Comparing the predicted structures of Klebsiella phage GH-K3 tail fiber protein (blue) and P. paraeruginosa PA7 CDS # 2309 (red). Note that the monomer form of Klebsiella phage depolymerase Depo32 (PDB #7VYV) was used as a reference to guide both structure predictions; this protein natively forms a homotrimer. Protein structure visualizations were done with PyMOL3.

### IPCD *Pseudomonas* genome dataset

Among the 1,271 *Pseudomonas* genomes investigated in this study, 1,387 phage tail proteins, stemming from 735 isolates, had a DePP score above 75%. However, 784 of those had no annotated Pfam domain and 496 of them had only a “Phage-tail_3” domain annotated. Nevertheless, among the remaining 107 phage tail proteins, ten harbored domains indicative of depolymerase activity (**Table 1**).

More precisely, eight had both a LysM motif (PF01476) and a transglycosylase SLT domain (PF01464) while two others had a PA2794-like exo-alpha-sialidase C-terminal domain (PF22432). These ten candidate PDPs, while not having the highest DePP score, have additional evidence that they contain domains capable of degrading cell wall polysaccharides and/or peptidoglycans. This allows to establish an *a priori* discovery rate of 1 candidate PDP per 100 isolates among the IPCD collection, strictly from a genomic data mining approach.

## Discussion

In this study, we have investigated the presence of prophage-borne depolymerases across 1,294 *Pseudomonas aeruginosa* genomes from the International Pseudomonas Consortium Database. We, therefore, developed a wrapper for Depolymerase-Predictor, called PDP-Miner, which annotates tail proteins *ab initio*, and then runs DePP exclusively on this subset while listing Pfam domains to further allow researchers to validate predictions made by DePP. We showed that this approach allowed to recover gene candidates whose amino acid translation had similar predicted 3D structures to known phage depolymerase structure.

To our knowledge, this is the first implementation of a tool that investigates phage depolymerase genes specifically within whole prokaryotic genomes, more precisely within the prophages they carry. The rationale for investigating whole genomes was that a prophage could infect its bacterial host at least once. Consequently, if it harbors depolymerase genes, then these gene products could degrade their host’s cell wall as well. If the infected host’s serotype is known, then a depolymerase from this cell’s prophages could theoretically degrade the cell wall of other strains harboring the same surface antigens. This allows us to predict a candidate depolymerase’s specificity for conspecific strains.

One limitation of this approach is whether the prophages have been rendered defective through neutral evolutionary forces over time, e.g. mutation and genetic drift (Bobay, Touchon and Rocha 2014). If a prophage is, by the result of these forces, unable to escape from its carrier, genes required for infection (including tail protein genes) will mostly drift due to the absence of selective pressure for resuming a lytic cycle. In any case, verifying a candidate depolymerase’s activity and specificity would require expressing and purifying the protein and then assessing its lytic ability *in vitro*.

PDP-Miner is also dependent on DePP’s intrinsic accuracy to predict depolymerase activity. As mentioned above, DePP outputs high probability scores even to non-phage proteins, suggesting that its learning model makes it recognize domains interacting with polysaccharides regardless of them being phage-associated or not (**Suppl. Table 1**). Among the IPCD genomes investigated, there were some instances of genes having high DePP scores (>90%) but without glycosyl hydrolase domains found. This could be attributable to stringent parameters passed to PfamScan, or to domains that have not yet been characterized. One example of this was the DUF1983 domain found in 4 of 5 of the type strains’ top scoring gene candidate. DUF stands for “Domain of Unknown Function” in Pfam nomenclature, however its association with both Klebsiella phage GH-K3 tail fiber protein and those top scoring candidates was notable, given the strong homology between all those (**Figure 3**).

As all machine learning tools, regardless of learning strategy, the quality of training data is paramount to the tool’s accuracy and recall capabilities (Gupta et al. 2021). DePP was trained on a 50/50 model including 50 experimentally verified phage depolymerase proteins and a random set of 50 proteins unrelated to phages (Magill and Skvortsov 2023). This dataset does not cover the diversity of phage depolymerases both from a taxonomic and structural point of view. Nevertheless, DePP allows for custom models to be built, which could be done when more experimental data will be available.

## Conclusion

PDP-Miner is a wrapper for Depolymerase-Predictor that augments its output with contextual information such as protein domains all while restricting search to phage tail proteins, thereby limiting false positives while mining whole genomes and their prophages. At a time when artificial intelligence and machine learning (AI/ML) are further and further involved in genome mining studies, it is important to contextualize outputs from these software as they tend to produce few output justifying their decisions. In this end, PDP-Miner provides an example of combining AI/ML with human oversight for interpreting bioinformatics results, all the while providing an informative workflow to identify gene candidates for the prospect of future antimicrobial compounds from phage/prophage products.

## Acknowledgements

We are indebted to Magill and Skvortsov for developing Depolymerase-Predictor (Magill and Skvortsov 2023), which served as a basis for the genome mining workflow discussed herein. We also thank anonymous reviewers for their critical advice and suggestions for this work. Work in the RCL laboratory was funded by The Canadian Institutes of Health Research, Genome Canada, Genome Québec and Ontario Genomics.

## Conflict of Interest Statement

The authors have no conflict of interest to declare.

## Notes

http:///www.github.com/jeffgauthier/pdpminer

### Competing Interest Statement

The authors have declared no competing interest.

## References

1. Antimicrobial Resistance Collaborators, ‘Global Burden of Bacterial Antimicrobial Resistance in 2019: A Systematic Analysis’, Lancet (London, England), 399/10325 (2022), 629–55

2. Barron, Madeline, ‘Phage Therapy: Past, Present and Future’, ASM.Org, 2022 <https://asm.org:443/Articles/2022/August/Phage-Therapy-Past,-Present-and-Future> [accessed 15 January 2025]

3. Bhattacharjee, Rahul, Nandi, Aditya, Sinha, Adrija, Kumar, Hrithik, Mitra, Disha, Mojumdar, Abhik, et al., ‘Phage-Tail-like Bacteriocins as a Biomedical Platform to Counter Anti-Microbial Resistant Pathogens’, Biomedicine & Pharmacotherapy, 155 (2022), 113720

4. Bobay, Louis-Marie, Touchon, Marie, and Rocha Eduardo P. C., ‘Pervasive Domestication of Defective Prophages by Bacteria’, Proceedings of the National Academy of Sciences, 111/33 (2014), 12127–32

5. Bouras, George, Nepal, Roshan, Houtak, Ghais, Psaltis Alkis James, Wormald, Peter-John, and Vreugde, Sarah, ‘Pharokka: A Fast Scalable Bacteriophage Annotation Tool’, Bioinformatics, 39/1 (2023), btac776

6. Coil, David, Jospin, Guillaume, and Darling Aaron E., ‘A5-Miseq: An Updated Pipeline to Assemble Microbial Genomes from Illumina MiSeq Data’, Bioinformatics (Oxford, England), 31/4 (2015), 587–89

7. Fleming, Alexander, ‘On the Antibacterial Action of Cultures of a Penicillium, with Special Reference to Their Use in the Isolation of B. Influenzæ’, British Journal of Experimental Pathology, 10/3 (1929), 226–36

8. Fong, Karen, Wong Catherine W.Y., Wang, Siyun, and Delaquis, Pascal, ‘How Broad Is Enough: The Host Range of Bacteriophages and Its Impact on the Agri-Food Sector’, PHAGE, 2/2 (2021), 83–91

9. Freschi, Luca, Jeukens, Julie, Kukavica-Ibrulj, Irena, Boyle, Brian, Dupont Marie-Josée, Laroche, Jérôme, et al., ‘Clinical Utilization of Genomics Data Produced by the International Pseudomonas Aeruginosa Consortium’, Frontiers in Microbiology, 6 (2015), 1036

10. Gupta, Nitin, Mujumdar, Shashank, Patel, Hima, Masuda, Satoshi, Panwar, Naveen, Bandyopadhyay, Sambaran, et al., ‘Data Quality for Machine Learning Tasks’, in Proceedings of the 27th ACM SIGKDD Conference on Knowledge Discovery & Data Mining, KDD ‘21 (New York, NY, USA, 2021), 4040–41 <10.1145/3447548.3470817> [accessed 15 January 2025]

11. Hutchings, Matthew I, Truman Andrew W, and Wilkinson, Barrie, ‘Antibiotics: Past, Present and Future’, Current Opinion in Microbiology, Antimicrobials, 51 (2019), 72–80

12. Ikuta, Kevin S, Swetschinski Lucien R, Robles Aguilar, Gisela, Sharara, Fablina, Mestrovic, Tomislav, Gray, Authia P, et al., ‘Global Mortality Associated with 33 Bacterial Pathogens in 2019: A Systematic Analysis for the Global Burden of Disease Study 2019’, The Lancet, 400/10369 (2022), 2221–48

13. Johnson, Genevieve, Banerjee, Swarnali, and Putonti, Catherine, ‘Diversity of Pseudomonas Aeruginosa Temperate Phages’, mSphere, 7/1 (2022), e0101521

14. Knecht, Leandra E., Veljkovic, Marjan, and Fieseler, Lars, ‘Diversity and Function of Phage Encoded Depolymerases’, Frontiers in Microbiology, 10 (2020) <https://www.frontiersin.org/journals/microbiology/articles/10.3389/fmicb.2019.02949/full > [accessed 15 January 2025]

15. Loc-Carrillo, Catherine, and Abedon Stephen T, ‘Pros and Cons of Phage Therapy’, Bacteriophage, 1/2 (2011), 111–14

16. Magill, Damian J., and Skvortsov Timofey A., ‘DePolymerase Predictor (DePP): A Machine Learning Tool for the Targeted Identification of Phage Depolymerases’, BMC Bioinformatics, 24/1 (2023), 208

17. Mistry, Jaina, Chuguransky, Sara, Williams, Lowri, Qureshi, Matloob, Salazar Gustavo A, Sonnhammer Erik L L, et al., ‘Pfam: The Protein Families Database in 2021’, Nucleic Acids Research, 49/D1 (2021), D412–19

18. Morin, Charles D., Déziel, Eric, Gauthier, Jeff, Levesque Roger C., and Lau Gee W., ‘An Organ System-Based Synopsis of Pseudomonas Aeruginosa Virulence’, Virulence, 12/1, 1469– 1507

19. NCBI, ‘CDD Conserved Protein Domain Family: CBM_5_12’, 2024 <https://www.ncbi.nlm.nih.gov/Structure/cdd/pfam02839> [accessed 15 January 2025]

20. Olszak, Tomasz, Shneider Mikhail M., Latka, Agnieszka, Maciejewska, Barbara, Browning, Christopher, Sycheva Lada V., et al., ‘The O-Specific Polysaccharide Lyase from the Phage LKA1 Tailspike Reduces Pseudomonas Virulence’, Scientific Reports, 7/1 (2017), 16302

21. Saha, Senjuti, Ojobor Chidozie D. Li; Annie Si Cong, Mackinnon, Erik, North Olesia I., Bondy-Denomy, Joseph, et al., ‘F-Type Pyocins Are Diverse Noncontractile Phage Tail-Like Weapons for Killing Pseudomonas Aeruginosa’, Journal of Bacteriology, 205/6, e00029–23

22. Taslem Mourosi, Jarin, Awe, Ayobami, Guo, Wenzheng, Batra, Himanshu, Ganesh, Harrish, Wu, Xiaorong, et al., ‘Understanding Bacteriophage Tail Fiber Interaction with Host Surface Receptor: The Key “Blueprint” for Reprogramming Phage Host Range’, International Journal of Molecular Sciences, 23/20 (2022), 12146

23. Wang, Honglan, Liu, Yannan, Bai, Changqing, and Leung, Sharon Shui Yee, ‘Translating Bacteriophage-Derived Depolymerases into Antibacterial Therapeutics: Challenges and Prospects’, Acta Pharmaceutica Sinica. B, 14/1 (2024), 155–69

24. Weimann, Aaron, Dinan Adam M., Ruis, Christopher, Bernut, Audrey, Pont Stéphane, Brown, Karen, et al., ‘Evolution and Host-Specific Adaptation of Pseudomonas Aeruginosa’, Science, 385/6704 (2024), eadi0908

25. WHO, ‘WHO Bacterial Priority Pathogens List, 2024: Bacterial Pathogens of Public Health Importance to Guide Research, Development and Strategies to Prevent and Control Antimicrobial Resistance’, 2024 <https://www.who.int/publications/i/item/9789240093461> [accessed 15 January 2025]

26. Winstanley, Craig, Langille Morgan G.I., Fothergill Joanne L., Kukavica-Ibrulj, Irena, Paradis-Bleau, Catherine, Sanschagrin, François, et al., ‘Newly Introduced Genomic Prophage Islands Are Critical Determinants of in Vivo Competitiveness in the Liverpool Epidemic Strain of Pseudomonas Aeruginosa’, Genome Research, 19/1 (2009), 12–23

27. Yang, Qimao, Le, Shuai, Zhu, Tongyu, and Wu, Nannan, ‘Regulations of Phage Therapy across the World’, Frontiers in Microbiology, 14 (2023) <https://www.frontiersin.org/journals/microbiology/articles/10.3389/fmicb.2023.1250848/full> [accessed 15 January 2025]

