## Supplementary material for "PDP-Miner: an AI/ML tool to detect prophage tail proteins with depolymerase domains across thousands of bacterial genomes": Suppl.

### Supplementary Figures

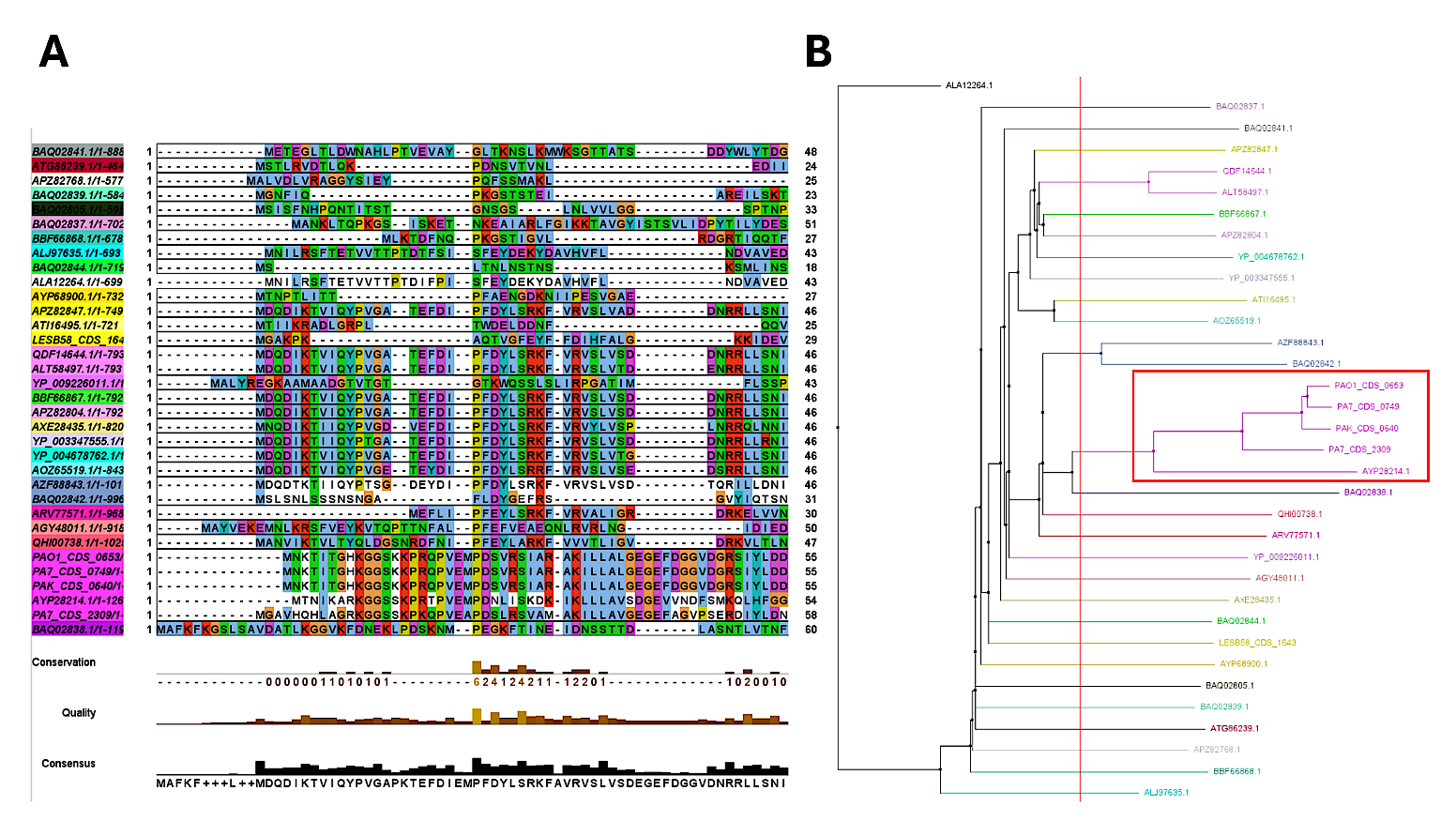

**Suppl. Figure 1.** **GH-K3 phage tail fiber protein (AYP28214.1) is the nearest neighbor to the top 5 PDPs. found by PDP-Miner.**  A : Multiple sequence alignment of the top 5 PDPs found in four *Pseudomonas* type strains, and DePP sequences used for model training. B: Neighbor-joining tree calculated from the alignment in A.

### Supplementary Tables

**Suppl. Table 1.** Top scoring genes according to Depolymerase-Predictor (DePP) when a whole-genome annotation scheme (Prokka) is used instead of a phage gene annotator (Pharokka).

All genes with a DePP probability score below 80% were discarded. Interestingly, all top scoring gene candidates were not phage-related, but were rather chromosome-encoded phospholipases, endonucleases and notably the autotransporter esterase EstA involved in rhamnolipid production. This finding implies that DePP is well trained at finding hydrolases (EC 3.1.x.x) with carbohydrate and/or lipid binding domains, and not necessarily phage tail proteins with depolymerase activity.

| **Genome** | **Locus** | **Abbr.** | **Description** | **EC number** | **COG number** | **Size (bp)** | **DePP probability** |
| --- | --- | --- | --- | --- | --- | --- | --- |
| Pa-LESB58 | CDS_05654 | estA | Esterase EstA | 3.1.1.1 | COG3240 | 1,941 | 91% |
|  | CDS_02946 | rhsC_3 | Putative deoxyribonuclease RhsC | 3.1.-.- | COG3209 | 2,637 | 89% |
|  | CDS_01756 | rhsC_1 | Putative deoxyribonuclease RhsC | 3.1.-.- | COG3209 | 2,658 | 88% |
|  | CDS_04611 | plcN_2 | Non-hemolytic phospholipase C | 3.1.4.3 | COG3511 | 2,193 | 86% |
|  | CDS_00042 | cdiA_1 | 16S rRNA endonuclease CdiA | 3.1.-.- | COG3210 | 10,611 | 85% |
|  | CDS_02943 | cdiA_2 | 16S rRNA endonuclease CdiA | 3.1.-.- | COG3210 | 16,938 | 82% |
|  | CDS_01806 | plcN_1 | Non-hemolytic phospholipase C | 3.1.4.3 | COG3511 | 2,079 | 82% |
| Pa-PA7 | CDS_05594 | estA | Esterase EstA | 3.1.1.1 | COG3240 | 1,941 | 91% |
|  | CDS_02669 | cdiA_4 | DNase CdiA | 3.1.-.- | NA | 2,871 | 86% |
|  | CDS_04466 | plcN_2 | Non-hemolytic phospholipase C | 3.1.4.3 | COG3511 | 2,193 | 84% |
|  | CDS_02665 | cdiA_3 | 16S rRNA endonuclease CdiA | 3.1.-.- | COG3210 | 10,692 | 83% |
|  | CDS_01719 | plcN_1 | Non-hemolytic phospholipase C | 3.1.4.3 | COG3511 | 2,079 | 81% |
| Pa-PAK | CDS_05381 | estA | Esterase EstA | 3.1.1.1 | COG3240 | 1,941 | 92% |
|  | CDS_02626 | rhsC_3 | Putative deoxyribonuclease RhsC | 3.1.-.- | COG3209 | 2,664 | 90% |
|  | CDS_01601 | rhsC_1 | Putative deoxyribonuclease RhsC | 3.1.-.- | COG3209 | 2,658 | 88% |
|  | CDS_04314 | plcN_2 | Non-hemolytic phospholipase C | 3.1.4.3 | COG3511 | 2,193 | 86% |
|  | CDS_00042 | cdiA | 16S rRNA endonuclease CdiA | 3.1.-.- | COG3210 | 10,512 | 86% |
|  | CDS_01650 | plcN_1 | Non-hemolytic phospholipase C | 3.1.4.3 | COG3511 | 2,079 | 83% |
| Pa-PAO1 | CDS_05276 | estA | Esterase EstA | 3.1.1.1 | COG3240 | 1,941 | 90% |
|  | CDS_00868 | plcN_1 | Non-hemolytic phospholipase C | 3.1.4.3 | COG3511 | 2,193 | 86% |
|  | CDS_00042 | cdiA_1 | 16S rRNA endonuclease CdiA | 3.1.-.- | COG3210 | 10,608 | 84% |
|  | CDS_03435 | plcN_2 | Non-hemolytic phospholipase C | 3.1.4.3 | COG3511 | 2,079 | 82% |
|  | CDS_02812 | hsdR | Type-1 restriction enzyme R protein | 3.1.21.3 | NA | 3,441 | 81% |
|  | CDS_02530 | cdiA_2 | 16S rRNA endonuclease CdiA | 3.1.-.- | COG3210 | 16,884 | 81% |
